# *Escherichia coli’s* physiology can turn membrane voltage dyes into actuators

**DOI:** 10.1101/607838

**Authors:** L Mancini, G Terradot, T Tian, Y Pu, Y Li, CJ Lo, F Bai, T Pilizota

## Abstract

The electrical membrane potential (*V*_*m*_) is one of the components of the electrochemical potential of protons across the biological membrane (proton motive force), which powers many vital cellular processes, and *V*_*m*_ also plays a role in signal transduction. Therefore, measuring it is of great interest, and over the years a variety of techniques has been developed for the purpose. In bacteria, given their small size, Nernstian membrane voltage probes are arguably the favourite strategy, and their cytoplasmic accumulation depends on *V*_*m*_ according to the Nernst equation. However, a careful calibration of Nernstian probes that takes into account the trade-offs between the ease with which the signal from the dye is observed, and the dyes’ interactions with cellular physiology, is rarely performed. Here we use a mathematical model to understand such trade-offs and, based on the knowledge gained, propose a general work-flow for the characterization of Nernstian dye candidates. We demonstrate the work-flow on the Thioflavin T dye in *Escherichia coli*, and identify conditions in which the dye turns from a *V*_*m*_ probe into an actuator.

**SIGNIFICANCE STATEMENT**

The phospholipid bilayer of a biological membrane is virtually impermeable to charged molecules. Much like in a rechargeable battery, cells harness this property to store an electrical potential that fuels life reactions but also transduces signals. Measuring this electrical potential, also referred to as membrane voltage, is therefore of great interest and a variety of techniques have been employed for the purpose, starting as early as the 1930s. For the case of bacteria, which are smaller in size and possess a stiffer cell wall, arguably the most popular approach to measuring membrane voltage are Nernstian probes that accumulate across the bacterial membrane according to the Nernst potential. The present study characterizes the undesired effects Nernstian probes can have on cell physiology, which can be crucial for the accurate interpretation of experimental results. Using mathematical modelling and experiments, the study provides a general, simple workflow to characterise and minimise these effects.

## INTRODUCTION

Living cells maintain an electric potential difference (*V*_*m*_) across the plasma membrane that acts like a capacitor, and is achieved by active transport of ions:

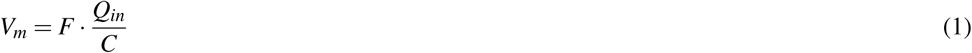

where *Q*_*in*_ is the intracellular charge (in mole), *C* the membrane capacitance and *F* the Faraday constant. Membrane potential stands at the basis of fundamental biological processes, such as signal transduction and energy production [Del Castillo and Katz, 1954, Mitchell, 1961]. For the latter, *V*_*m*_ adds up to the chemical potential of protons, arising from their concentration difference across the membrane, to result in the proton electrochemical gradient, so called proton motive force (PMF). The PMF drives numerous cellular processes, most notably the production of ATP [Mitchell, 1961], import of nutrients or osmolites [Bradbeer, 1993, Jahreis et al., 2008, Ramos and Kaback, 1977, Wood, 2015] rotation of the bacterial flagellar motor [Sowa and Berry, 2008], and it is necessary for cell division [Strahl and Hamoen, 2010].

The notion that *V*_*m*_ lies at the very basis of life motivated decades long efforts to measure it. The first technique dates to 1939 and relies on the mechanical insertion of microelectrodes into squid giant axons [Hodgkin and Huxley, 1939]. The method was later applied to neurons [Ling and Gerard, 1949], and led to the development of the patch-clamp technique, which advanced the understanding of neuron signal transduction [Neher and Sakmann, 1976, Sakmann and Neher, 1984]. However, applicability of microelectrodes for the measurement of bacterial *V*_*m*_ is limited, owing to the small size of the organisms and the presence of the cell wall [Martinac et al., 1987, 2013]. Some of the subsequently developed methods overcome such limits with the use of molecular sensors [Felle et al., 1980], grouped in two categories: conformational-change-based sensors and Nernstian sensors. The former are static molecules or proteins that sit inside the membrane, or in its close proximity, and change conformation or electron distribution in response to changes in *V*_*m*_, which in turn affect the optical properties of the chromophores thus transducing signal [Fluhler et al., 1985, Kralj et al., 2011, Tsutsui et al., 2008]. Here we focus on the latter, the Nernstian sensors, and on the parameter range in which they serve as *V*_*m*_ indicators, using *Escherichia coli* as the model organism.

Nernstian sensors are charged molecules that can diffuse across the biological membranes and distribute according to the Nernst equation:

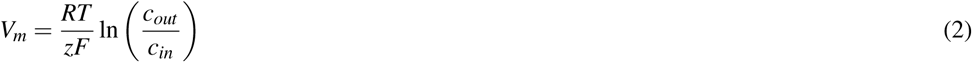

where *R, T*, *z, F, c*_*out*_, *c*_*in*_ denote respectively gas constant, temperature, valence of the charged molecule, Faraday’s constant, external and internal concentration of the charged molecule. For a measurement to be attained, these molecules need to emit a signal that is a proxy for their number. Therefore, Nernstian *V*_*m*_ dyes are usually radio-labeled or fluorescent molecules [Felle et al., 1980, Sims et al., 1974], and *V*_*m*_ is calculated from equation (2) by measuring the cytoplasmic (*c*_*in*_) and the external dye concentration (*c*_*out*_ [Lo et al., 2007].

However, Nernstian dyes are used in complex biological systems and a number of factors can be responsible for an incomplete adherence to a fully Nernstian behavior. In Fig. 1 we give a cartoon representation of the trade-offs imposed on a Nernstian dye by plotting the dye intensity inside *E. coli’s* cytoplasm against the time. The chosen dye concentration should be such that the signal is sufficiently above the background (Δ*I* is sufficiently large). Yet, with increasing dye concentration, cell’s *V*_*m*_ is more likely to be affected by the dye. This caveat is inherent to positively charged dyes as these directly lower *V*_*m*_ and more so at higher concentrations [Kashket, 1985]. The first requirement for a Nernstian dye is, thus, existence of a range of concentrations that give sufficient signal without extensively affecting the *V*_*m*_. Likewise, cellular processes should not interfere with the Nernstian behavior of the dye, for example by actively importing or exporting it. Instead, the dye should be able to diffuse across the membrane and its diffusion constant will determine the time it takes for the dye to equilibrate across the membrane in agreement with equation (2) (*τ*_*eq*_ in Fig. 1). All phenomena that occur quicker than *τ*_*eq*_ are beyond the dye’s temporal resolution, and likewise, all the measurements taken before *τ*_*eq*_ do not faithfully report *V*_*m*_. Lastly, for quantitative measurements, Nernstian dyes should exhibit a well defined and constant correlation between the concentration and the signal, e.g. the dye should not self-quench at any point [te Winkel et al., 2016] or undergo signal enhancements.

**Figure 1.**
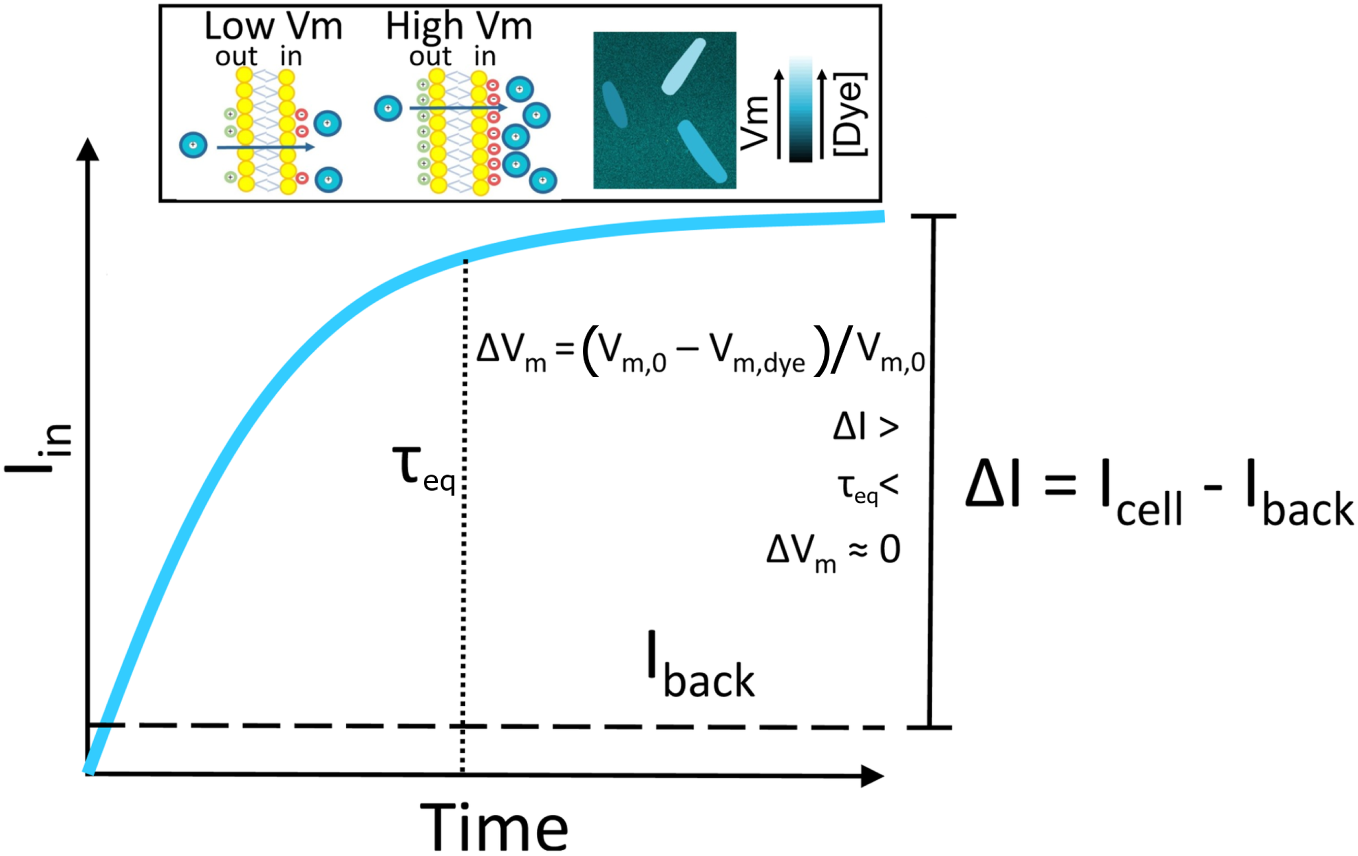
A schematic plot of a Nernstian dye equilibration curve. Equilibration time, *τ*_*eq*_, is defined as the time at which the internalized dye *I*_*in*_ reaches the 90% of its final value. *V*_*m*,0_ and *V*_*m,dye*_ indicate the membrane potential before and after the addition of the dye, respectively. Inset: cartoon showing the mechanism of accumulation of cationic dyes, which accumulate more in cells with a highly negative *V*_*m*_.

**Figure 2.**
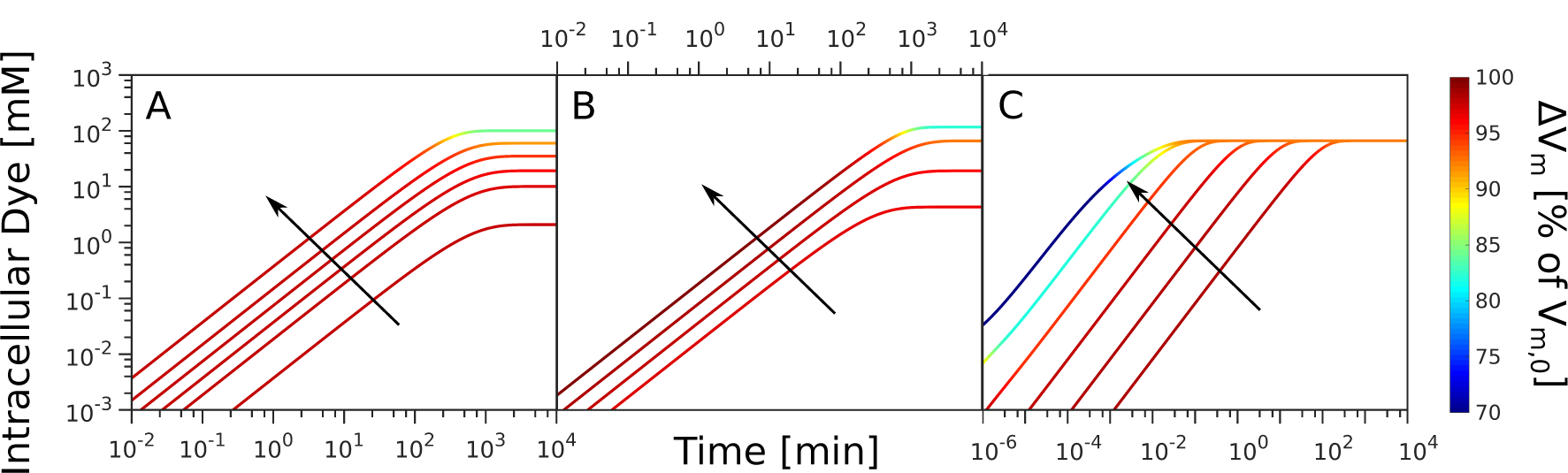
Computational data describing the parameter landscape associated with cationic dye usage as Nernstian sensors. (A) *V*_*m*,0_ = –140 mV, *Y*_*i*_ = 150 mM, *I*_*Na*_ = 210 mV. Intracellular Dye concentration as a function of time, for extracellular dye concentrations 10, 50, 100, 200, 400 and 1000 *µ*M. The arrow indicated increasing [*Dye*]_*out*_. (B) Extracellular dye concentration of 100 *µ*M, *Y*_*i*_ = 150 mM, Δ*G*_*E*_ = –210 mV. Intracellular dye concentration as a function of time, for different *V*_*m*,0_: –220, –180, –140 and –100 mV. The arrow indicates increasing absolute value of the *V*_*m*,0_. (C) *V*_*m*,0_ = –180 mV, *Y*_*i*_ = 150 mM, Δ*G*_*E*_ = –210 mV, Extracellular dye at 100 *µ*M. Intracellular dye concentration as a function of time, for different apparent permeabilities of the membrane to the dye: 10^−12^, 10^−10.8^, 10^−9.6^, 10^−8.4^, 10^−7.2^, 10^−6^ meter per second. The arrow indicates increasing permeability.

To summarize, to be used as a Nernstian sensor, a cationic dye should: (i) give a sufficiently high signal without affecting cell’s *V*_*m*_; (ii) diffuse through the membrane with *τ*_*eq*_ on the order of minutes; (iii) stay inert, despite being charged, and not form bonds or in any way interact with cell; (iv) have constant signal per molecule. Yet, when using such dyes these requirements are rarely assessed in a systematic manner before measurements commence. In this work we identify a work-flow that should be adopted in order to identify the parameter range in which Nernstain dyes act as sensors, rather than actuators. We start with a mathematical model that helps us understand relationships and define trade-offs between dye working concentration and signal intensity, equilibration time and *V*_*m*_ perturbation. We then show how the identified work-flow can be used to benchmark new Nernstian dyes by characterizing the recently reported dye Thioflavin T (ThT) used in *Bacillus subtilis* [Prindle et al., 2015], for use in *E. coli*. We describe the physiological range in which ThT enables *V*_*m*_ sensing in *E. coli*, and, in the range where we find it turns into an actuator, we investigate the mechanistic reasons. Our work-flow can be applied to the characterization of other Nernstian dyes and provide novel insights for the established ones.

## METHODS

### Bacterial strains

All experiments where no mutation is explicitly indicated were carried out in the MG1655 strain. For the BFM speed assay we used MG1655 carrying the FliCsticky mutation from [Krasnopeeva et al., 2018]. Δ*tolC* mutants were obtained from the Keio collection [Baba et al., 2006]. Kanamycin resistance of the Keio deletion strain was removed via one-step inactivation with the plasmid pCP20 [Datsenko and Wanner, 2000]. Kanamycin resistance inactivation and elimination of the pCP20 plasmid were confirmed via Kanamycin, Chloramphenicol and Ampicillin sensitivity tests. Both the strain carrying the Δ*tolC* mutation and MG1655 wild-type were transformed with plasmid pTP20-mKate2 (Fig. SI9) for cytoplasmic volume measurements. pTP20-mKate2 contains the red fluorescent protein mKate2 and the ribosomal binding site (RBS) of mCherry. The plasmid was constructed as follows: the backbone from pWR20 [Pilizota and Shaevitz, 2012] and the sequence containing the RBS of mCherry and mKate2 were PCR amplified. The products were purified, cleaved with the restriction enzymes AvrII and NotI (NEB, UK) and ligated using T4 DNA ligase (Promega, UK). Chemically competent cells were transformed with the ligation mixes and transformants were confirmed by colony PCR and subsequently sequenced. A map of the plasmid and the primers are given in SI (Fig. SI9, and Table SI1). All the strains used in the study are summarized in the Table SI2.

### Bacterial growth conditions

Cells for fluorescence microscopy were grown from an overnight culture by diluting it 1:80 times in LB (0.5% Yeast Extract, 1% Bacto tryptone, 0.5% NaCl). The culture was shaken at 220 rpm at 37°C and harvested at *OD*_600_=0.3-0.5. Upon harvest we washed the cells into fresh LB or MM9 + glucose medium (50mM Na_2_HPO_4_, 25mM NaH_2_PO_4_, 8.5mM NaCl, 18.7mM NH4Cl, 0.1mM CaCl_2_, 1mM KCl, 2mM MgSO_4_, 1x MEM essential amino acids (Gibco, UK) and 0.3% glucose). For the simultaneous BFM speed and ThT fluorescence measurements cells were grown from an overnight culture by diluting it 1:80 times in TB (1% Bacto tryptone, 0.5% NaCl) at 200 rpm and 30°C. Cells were harvested at *OD*_600_=0.8 as before [Rosko et al., 2017] and washed to fresh MM9 via centrifugation. Growth curves were obtained in a Spectrostar Omega microplate reader (BMG, Germany) using a flat-bottom 96-well plate that was covered with a lid during the experiments (Costar, UK). Each well contained 200*µ*l of growth media, either MM9 + glucose or MM9 + glycerol (50mM Na_2_HPO_4_, 25mM NaH_2_PO_4_, 8.5mM NaCl, 18.7mM NH4Cl, 0.1mM CaCl_2_, 1mM KCl, 2mM MgSO_4_, 1x MEM essential aminoacids (Gibco, UK) and 0.3% glycerol), and was inoculated with 2*µ*l (1:100 dilution) of an overnight culture. Plates were grown at 37°C with 300 rpm shaking (double orbital mode). ThT (Acros organics, USA) solutions were prepared from a 10 mM stock of ThT in water made at least monthly and stored at 4°C in the dark.

### Fluorescence microscopy

Imaging was carried out in a custom-built microscope with a 100x oil immersion objective lens (Nikon, Japan), Neutral White LED as a source of illumination (Cairn Research Ltd, UK) and images were taken with an iXon Ultra 897 EMCCD camera (Andor, UK) [Krasnopeeva, 2018, Rosko, 2017]. ThT fluorescence was measured with ZET436/20x and ET525/40m, and mKate2 and PI fluorescence with ET577/25x and ET632/60m (Chroma Technology, USA) excitation and emission filters, respectively. Exposure time was 50 ms and Andor camera gain 25. We note that ThT undergoes a spectral shift and intensity increase when highly concentrated or when spatially constricted, either by binding to amyloid fibrils or by viscosity [Maskevich et al., 2015, 2007, Sulatskaya et al., 2017]. Our choice of filters aims at minimizing these effects and the damage that shorter wavelengths cause to *E. coli* [Vermeulen et al., 2008]. Cells were imaged in a custom-built flow-cell (Fig. SI10, [Krasnopeeva et al., 2018]), and attached to the coverslip surface as before [Krasnopeeva et al., 2018, Rosko et al., 2017]. Briefly, 1% Poly-L-Lysine (Sigma, UK) is flushed through the flow cell and washed with 3-5 ml of growth media after 10 s. Polystyrene particles (beads) with a diameter of 1 *µ*m (Bangs Laboratories, USA), were delivered into the flow-cell and allowed to attach to the coverslip surface. After 10 min unattached beads were flushed away with 1-2 ml of growth media. Next, 200 *µ*l of cells were delivered to the flow-cell and allowed to attach for 10-30 min, after which the unattached cells were removed with 1 ml of growth medium. 10 *µ*M ThT in growth media was delivered with a peristaltic pump (Fusion 400, Chemyx, USA) using 50 *µ*l/min flow rate while imaging. We deliver 5 *µ*M of PI stain (MP Biomedicals, USA) in the same way. 5mM PI stock solution (in water) was stored at 4°C in the dark. Images were stabilized in x, y and z position using a bead attached to the cover-slip and back-focal-plane interferometry [Buda et al., 2016, Pilizota and Shaevitz, 2012].

### Motor speed measurements

Single motor speeds were measured as before [Krasnopeeva et al., 2018, Rosko et al., 2017]. Briefly, we sheared flagellar filaments by passing them through two syringes with narrow-gauge needles (26 gauge) connected by plastic tubing. The cell attachment protocol was as above, except 0.5 *µ*m beads (Polysciences, USA) were delivered after cell attachment allowing them to attach to filament stubs. Motor speed was measured during continuous flow that delivered MM9 + glucose medium supplemented with 10 *µ*M ThT. Back-focal-plane interferometry setup and recording conditions are as before [Rosko et al., 2017].

### Data analysis

#### Motor speed traces

Raw traces of the position of the bead attached to the filament stub were analyzed by a moving-window discrete Fourier transform as in [Rosko et al., 2017]. From the obtained motor speed traces DC frequency (50 Hz) was removed, speeds lower then 5 Hz ignored, and subsequently a median filter (window size 11) was applied [Krasnopeeva et al., 2018]. We note that we use a wild type strain for which the BFM can change rotational direction, which appears as a negative speed after application of the moving-window Fourier transform. However, for the purpose of PMF measurements these short intervals can be disregarded, and we only show the speed values above 0 Hz.

#### Fluorescence images

The image analysis was carried out with a custom written software. From fluorescence images, rectangles containing ‘flat’ cells, i.e. cells that are uniformly attached to the coverslip surface, as well as background rectangles within each cell-containing rectangle, were manually selected [Buda et al., 2016, Pilizota and Shaevitz, 2012]. The edge of the cell was identified within the cell-containing rectangle by applying a global threshold via the Otsu’s method [Otsu, 1979]. To total cells’ intensity values were obtained by summing up and averaging pixel belonging to the cells. Values obtained from the background rectangles at the time points when ThT was loaded in the channel but cells had not taken it up yet, were subtracted from the cell intensity values. The beads used for image stabilisation stain easily with ThT, and were used as a point of reference for dye entry (which in our case occurred 7 to 10 min from the start of imaging). We show fluorescence intensity traces that start at the point of ThT entry, but note that cells were exposed to fluorescence illumination in the 7-10 min interval before. For the low fluorescence values characteristic of the early stages of dye equilibration, our script fails to identify cells, in which case we linearly interpolate values between two closest events of successful cell identification. Cell area was measured from intensity profiles, by normalizing them and counting the pixels above 30% of maximum intensity as described previously [Buda et al., 2016, Pilizota and Shaevitz, 2012]

#### Plate reader data

Individual growth curves were analysed with the software deODorizer from [Swain et al., 2016]. To extract the maximum growth rate, 3 or more repeats in the same condition were aligned by the chosen OD value (usually OD *∼*0.4) using the growth curve that reached it first (in the given condition). The maximum growth rates given in Fig. 3C were normalized by the maximum growth rate in [Dye]_*out*_ =0 condition.

**Figure 3.**
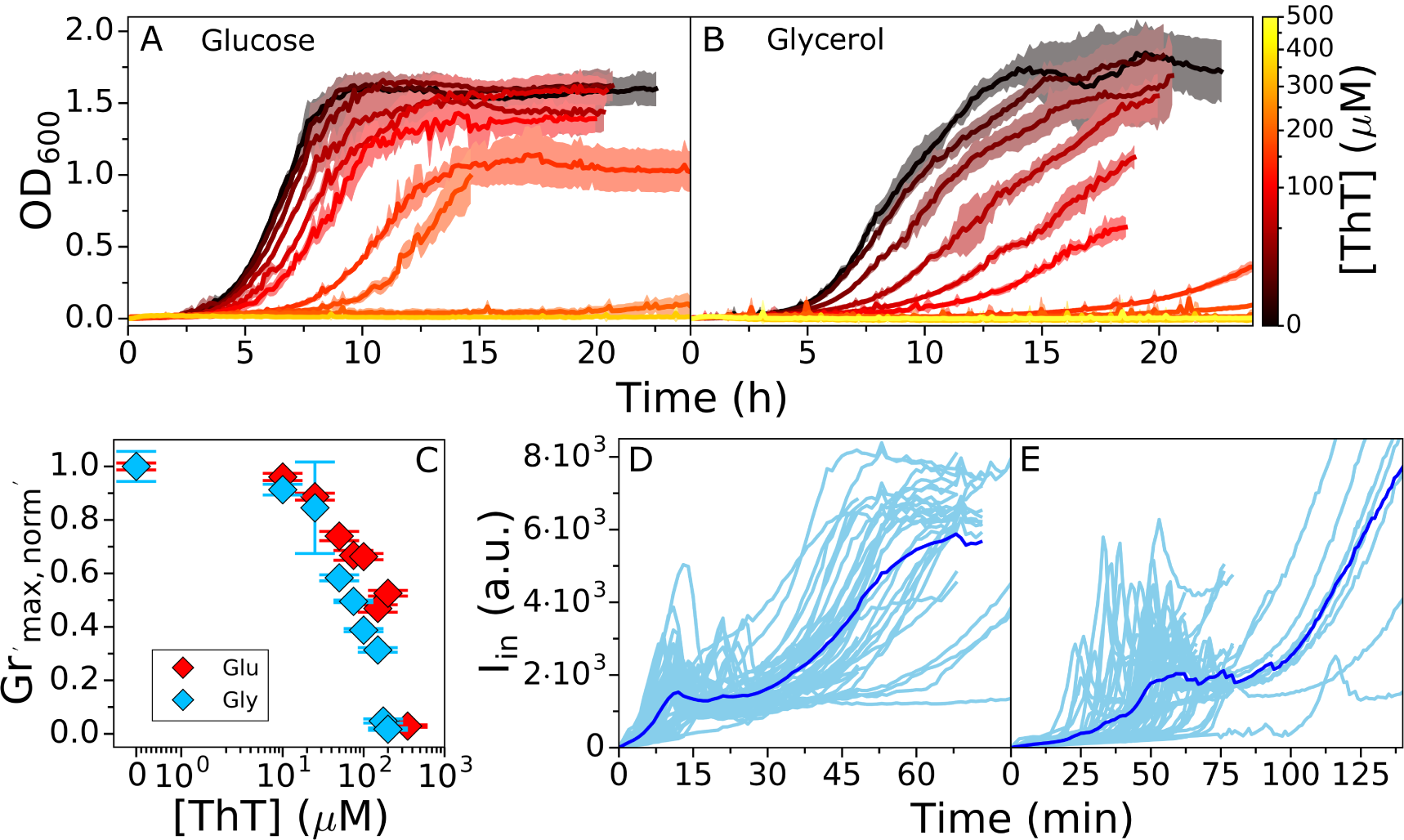
*E. coli* growth in the presence of ThT. *E. coli* growing in MM9 media supplemented with glucose or (B) glycerol at increasing ThT concentration (colourmap). ThT concentrations in (A) are 10, 25, 50, 75, 100, 200, 350 *µ*M and in (B) 10, 25, 50, 75, 100, 150, 175, 200 *µ*M). The error bars are standard deviations. (C) Maximum growth rates from (A) and (B) for each ThT concentration are given in red and blue respectively. Each condition was done at least in triplicate and error bars are the standard deviation. (D) *I*_*in*_ against time in LB and in (E) MM9 + glucose media. Individual cells are shown in cyan (45 in (D) and 52 in (E) from at least 9 independent experiments), and the average trace is shown in blue. The imaging conditions and *I*_*ex*_ = 10*µM* are the same for (E) and (D).

## RESULTS

### Mathematical model of Nernstian dye’s behavior defines its working parameter range

To predict and understand the mutual effects of dye concentration and cell physiology we turn to a mathematical model. We assume that the cytoplasmic and extracellular liquids are electrical conductors separated by the membrane, which we treat as a parallel-plate capacitor (equation 1) [Grabe and Oster, 2001]. We account for four types of charge carriers and assume that all are monovalent to simplify the model without altering the results with respect to *V*_*m*_ dye behavior: *(i)* negatively charged molecules to which the membrane is close to non-permeable denoted *Y*, *(ii)* cationic species actively pumped outward denoted *C*^+^, *(iii)* anionic species, which equilibrates across the membrane *A*^−^ and *(iv)* cationic species that equilibrate across the membrane playing the part of a cationic dye. Thus *Q*_*in*_ is:

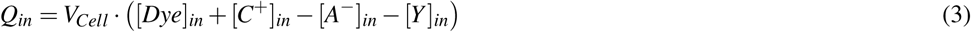

where *V*_*cell*_ is the intracellular volume. The extracellular concentrations and [*Y*]_*in*_ are constants set by the initial conditions (we assume that the cell does not affect the ionic composition of its environment and we treat [*Y*]_*in*_ as unable to cross the membrane). We also note that [*Dye*]_*in*_ and [*Dye*]_*out*_ are experimentally determined from fluorescence intensity signal, thus when presenting experimental results we use *I*_*in*_ and *I*_*out*_ instead.

The charge separation, and so *V*_*m*_, is achieved in two ways. First, by pumping 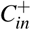 outwards and thus creating an overall negatively charged intracellular environment, and second by maintaining [*Y*]_*in*_ (an example of [*Y*]_*in*_ is glutamate). We set the free-energy for the outward pumping of *C*^+^ against its electrochemical gradient to be a constant, and label it Δ*G*_*E*_ (where Δ*G*_*E*_ *<* 0). For example, in the case of a proton:ion antiporter with 1:1 exchange stoichiometry the free energy is the PMF itself, and for a similar antiporter with 2:1 proton:ion stoichiometry it is 2 *×* PMF.

The rate at which *C*^+^ is pumped out of the cell is 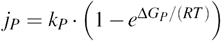, where *k*_*P*_ is a function that describes the specifics of the transport mechanism by a given pump, here, we keep it a constant. Δ*G*_*P*_ depends on the electrochemical potential of the pumped cation 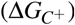 and Δ*G*_*E*_. The chosen functional dependency of *j*_*P*_ gives the simplest pump kinetics, sufficient for our purpose, which can be expanded to include more complex pumping scenarios Keener and Sneyd [2009]. Therefore, the rate of pumping (positive flux means *C*^+^ is extruded) depends on the intracellular ionic composition via *V*_*m*_ and [*C*^+^]_*in*_:

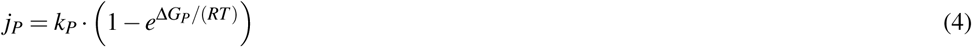

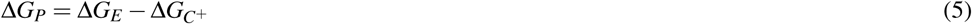

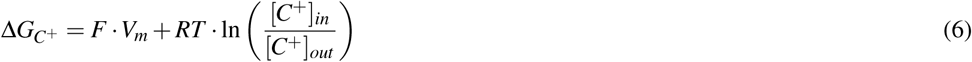

Note that in order for the pump to move *C*^+^ outward *j*_*P*_ *>* 0, and consequently 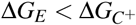, i.e. the free-energy providing reaction has to be able to overcome the electrochemical gradient of the *C*^+^.

Finally, the dye, the anion and the cation leak through the membrane (positive flux means *x* is moved inward) at the rate:

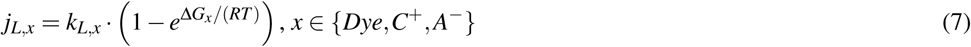

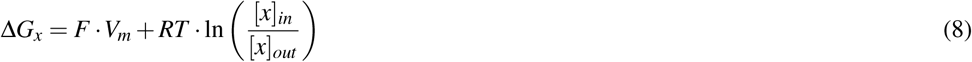

Similarly to *k*_*P*_, *k*_*L,x*_ is a function whose shape depends on the mechanisms by which an ion leaks across the *E. coli* membrane, which in turn depends on the electrostatic potential at a position *z* within the membrane, *V* (*z*). To the best of our knowledge, *V* (*z*), and consequently *dV* (*z*)*/dz*, are not known for *E. coli*. We chose Eyring’s model that has been verified for cationic leakage across the mitochondrial membrane [Garlid and Paucek, 2003], and which assumes *V* (*z*) abruptly changes in the middle of the lipid bilayer, such that *dV* (*z*)*/dz* = 0 everywhere but at the geometrical middle of the membrane where *dV* (*z*)*/dz* = *V*_*m*_ [Garlid et al., 1989]. We then have:

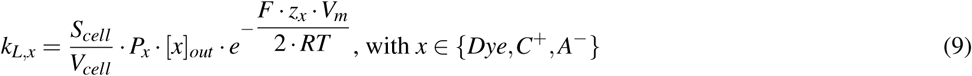

where *S*_*cell*_ denotes the cell’s surface area and *P*_*x*_ the permeability of the membrane for *x ∈*{*Dye,C*^+^, *A*^−^} At steady-state *Dye* and *A*^−^ equilibrate across the membrane according to Nernst equation (*d*[*Dye*]_*in*_*/dt* = *j*_*L,Dye*_ = 0 *⇔*Δ*G*_*Dye*_ = 0, leading to (2)), whereas for the monovalent cation 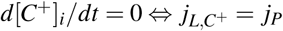. Next we introduce a new variable (“*pump-leak ratio*”) defined as:

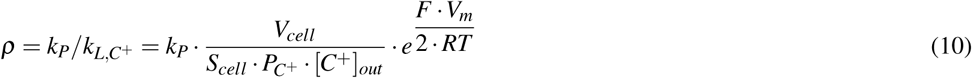

and re-write the steady-state condition for *C*^+^ as:

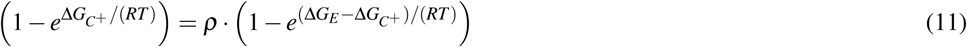

Given a certain extracellular composition ([*Dye*]_*out*_, [*C*^+^]_*out*_ and [*A*^−^]_*out*_), and taking into account that [*Dye*]_*in*_ and [*A*^−^]_*in*_ are defined by the Nernst equation at steady-state, equation (11) gives us a unique solution for steady-state *V*_*m*_ for a set of {[*Y*]_*in*_, ***ρ***, Δ*G*_*E*_} values.

We note from Eq. (11) that changing the functional dependency of *k*_*L,x*_ or *k*_*P*_ does not affect how the steady-state *V*_*m*_ depends on *ρ*. However, the steady-state potential after addition of the dye relative to the steady-state potential in absence of the dye (Δ*V*_*m*_ = (*V*_*m*,0_–*V*_*m,Dye*_)*/V*_*m*,0_) does. Similarly, the dynamics of dye equilibration are dependent on the functional dependency of *k*_*L,x*_, *k*_*P*_, *e.g.* assuming Goldman–Hodgkin–Katz flux equation Goldman [1943] instead of Erying’s model for *k*_*L,x*_ will give a slightly different dye equilibration profile (Fig. SI1). However, such changes will not influence the conclusions we reach based on our model predictions, because we are interested in the changes of the intracellular dye concentration dynamics at, e.g. different extracellular dye concentrations, *V*_*m*,0_ or *P*_*Dye*_. These partial derivatives of the intracellular dye concentration are invariant to the choice of *k*_*L,x*_ and *k*_*P*_.

Having constructed the model, we obtain the computational data in Fig. 2 in two steps. In the first step we allow the ODE system described by Eqs. SI 20, 21 to reach the steady-state (*V*_*m*,0_) for a 3-D grid of {[*Y*]_*in*_, *ρ*, Δ*G*_*E*_}. We note that in this step we do not need to specify ions’ permeabilities nor the rate function for leakage *k*_*L,x*_, because we define the values of *ρ*, which is the ratio of the two. We then use the obtained *V*_*m*,0_ as the initial condition for the second step of the numerical experiment, which requires us to specify: *(i)* the rate function for leakage (Eq. (9)), *(ii)* the permeability of the membrane to the dye *P*_*Dye*_ and *(iii)* the concentration of the dye in the extracellular space [*Dye*]_*out*_.

As increasing the [*Dye*] gives better signal-to-noise ratio, we first look at the dye equilibration profile ([*Dye*]_*in*_) for fixed *V*_*m*,0_, [*Y*]_*i*_ = 150 mM, Δ*G*_*E*_ = –210 mV and at different external dye concentrations ([*Dye*]_*out*_). As expected, Fig. 2A shows that increasing [*Dye*]_*out*_ improves the signal-to-noise ratio and shortens *τ*_*eq*_, but at the same time increasingly depolarizes the membrane. The extent by which Δ*V*_*m*_ drops does not solely depend on the [*Dye*]_*out*_, but also on the initial *V*_*m*,0_. Fig. 2B shows dye equilibration profile for a fixed [*Dye*]_*out*_, but for different *V*_*m*,0_ indicating that highly polarized cells are more susceptible to *V*_*m*_ loss. Apart from the value of *V*_*m*,0_, Δ*V*_*m*_ will also depend on the charged permeable and non-permeable species that are generating it, as shown in Fig. SI2. If a given *V*_*m*,0_ value is generated in the presence of a higher concentration of charged, impermeable intracellular species or at a higher energetic cost, Δ*V*_*m*_ will increase for the same [*Dye*]_*out*_. Thus, the extent to which a given [*Dye*] becomes and actuator and affects the Δ*V*_*m*_ is context dependant. Furthermore, while increasing [*Dye*]_*out*_ shortens *τ*_*eq*_ this is only the case when *V*_*m*,0_ is affected, as seen in Fig. SI3. Lastly, we look at the dye equilibration profile for different permeabilities of the membrane to the dye (*P*_*Dye*_) in Fig. 2C, and show that for higher *P*_*Dye*_ same concentration of the dye lowers *V*_*m*,0_ more. Fig. SI4 shows *τ*_*eq*_ as a function of *P*_*Dye*_ for different *V*_*m*,0_.

### The working concentration of Nernstain dye Thioflavin T for *E. coli* is in *µ*m range

Guided by the model predictions we devise an experimental work-flow for assessing the parametric range in which a candidate cationic dye behaves like a Nernstian sensor, and choose Thioflavin T (ThT) for the purpose. ThT has recently been used as a *V*_*m*_ dye in *B. subtilis* [Prindle et al., 2015], but has not been characterized for use in *E. coli*. We start by identifying the working concentration that gives sufficiently large signal, yet minimizes the membrane voltage perturbation, Δ*V*_*m*_, and use the growth rate as a proxy for affected Δ*V*_*m*_. Fig. 3A and B show *E. coli* growth curves in MM9 media supplemented with glucose or glycerol respectively (see *Materials and Methods* for detailed media composition), and in the presence of a range of ThT concentrations. Fig. 3C shows growth rates plotted against the ThT concentration for both media, demonstrating that 10 *µ*M ThT or less does not significantly affect the growth rate in either media. We call this concentration MNC (Maximum Non-inhibitory Concentration). However, the growth rate reduction observed for higher ThT concentrations is media dependent (Fig. 3C). The result is consistent with the finding of our model that the effect of the dye on cell’s physiology is environment dependent.

We next check that the highest ThT concentration, which does not affect the growth rate (MNC), 10 *µ*M, gives sufficiently high signal-to-noise ratio by observing the dye equilibration in different media. We note that if sufficient Δ*I* is achieved with 10 *µ*M ThT we would further check that Δ*V*_*m*_ *<* 1% by measuring the *τ*_*eq*_ with both 10 *µ*M and a lower dye concentration. If Δ*V*_*m*_ *<* 1%, we expect *τ*_*eq*_ not to change based on the results of our model (Fig. 2A). Fig. 3D shows *I*_*in*_ in time in LB and Fig. 3E the same in MM9 media supplemented with glucose. In both cases fresh media with ThT is continuously supplied using a customized flow-cell (see *Materials and Methods*), and in both cases Δ*I* is sufficiently high. However, observed profiles are different from expected (Fig. 1), and show a characteristic initial peak and a final plateau (SI Video 1). We reasoned that the peak could either be a real fluctuation in *V*_*m*_ or it could indicate an unknown dye export mechanism.

### Multi-drug efflux pumps influence ThT accumulation in *E. coli*

To determine whether the observed peak in *I*_*in*_ is due to active export of the dye we first check that in *E. coli* ThT is not a multi-drug-efflux pump substrate. We are motivated by previous reports that show that dyes such as ethidium bromide and Nile red are substrates of pumps belonging to the five bacterial structural families: ABC (ATP-binding cassette), RND (resistance/nodulation/division), MATE (multidrug and toxic compound extrusion), MFS (major facilitator superfamily) and SMR (small multidrug resistance) [Alvarez-Ortega et al., 2013, Bay et al., 2008, Kuroda and Tsuchiya, 2009, Lubelski et al., 2007, Nikaido and Takatsuka, 2009]. Fig. 4A shows dye equilibration curves in a wild type (WT) strain compared to the strain bearing a deletion of TolC, which is a gene encoding for an outer membrane protein (OMP) that is a ubiquitous component of multi-drug efflux pumps [Anes et al., 2015]. The *I*_*in*_ peak in the deletion mutant did not disappear, instead the intensity level of the peak was even higher, suggesting the pump is involved in the dye export, but also that the qualitative difference between the expected and the observed equilibration curve shape is not due to dye export. Interestingly, in the mutant, the peak also occurred earlier in time during the loading and with less cell-to-cell variability. We next tested the effect of the ThT dye on the Δ*TolC* mutant growth rates, Fig. 4B. We found that at the MNC for the WT, the mutants’ growth was inhibited over the course of our experiment. Taken together the results indicate that ThT is likely pumped out of the cells by one or more types of TolC-dependent efflux pumps, thereby preventing its use for quantitative *V*_*m*_ measurements in WT *E. coli*.

**Figure 4.**
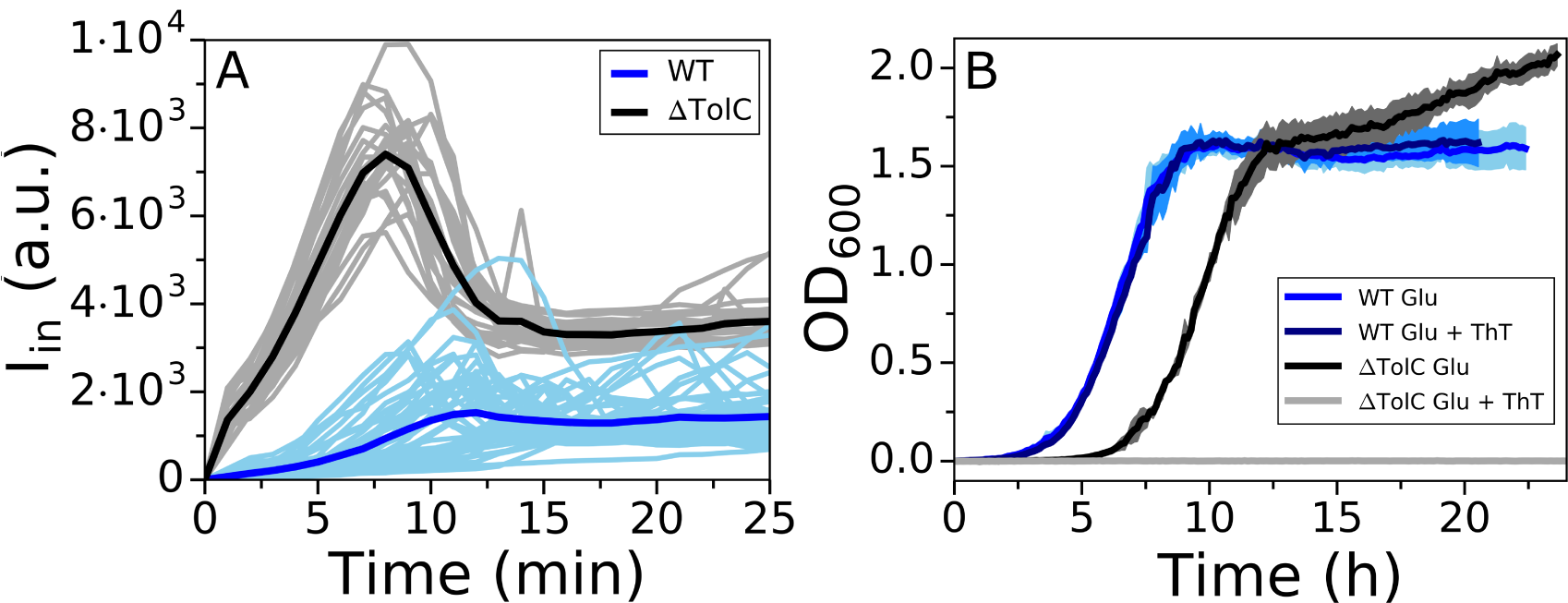
Comparison of WT and ΔTolC mutant response to ThT. (A) *I*_*in*_ versus time for the WT (cyan) and ΔTolC (gray) loaded in LB. WT traces are reproduced from Fig. 3D and ΔTolC traces were obtained from 5 independent experiments to give 23 single cell traces. Averaged traces for the WT and ΔTolC are given in blue and black respectively. (B) Growth curves of WT and ΔTolC in MM9 + glucose media are given in blue (reproduced from Fig. 3A) and black, respectively. Growth curves in the same media, but in the presence of 10 *µ*M ThT, are given in cyan (WT) and gray (ΔTolC). The shaded areas show standard deviation and cyan and blue growth curves for the WT overlap.

### Changing the membrane permeability during ThT loading can lead to loss of *V*_*m*_

We next tested our second hypothesis that the *I*_*in*_ peak is due to a decrease in *V*_*m*_. To this end, we performed measurements of bacterial flagellar motor (BFM) speed [Krasnopeeva et al., 2018] during ThT loading. BFM is a rotary molecular motor roughly 50 nm in size that enables bacterial swimming [Sowa and Berry, 2008] via PMF driven rotation [Fung and Berg, 1995, Manson et al., 1980, Matsuura et al., 1977, Meister and Berg, 1987]. The motor speed (*ω*) varies linearly with PMF [Fung and Berg, 1995, Gabel and Berg, 2003], which enables its use as a PMF, and when *pH*_*in*_ = *pH*_*out*_, as a *V*_*m*_ indicator as well [Krasnopeeva et al., 2018]. In our conditions, *pH*_*out*_ is 7 and *pH*_*in*_ is 7.86 (Fig. SI5) making the contribution to the PMF from ΔpH *∼*50 mV. Thus, even if during our experiment ΔpH goes to 0, we can learn about the *V*_*m*_ behaviour from the PMF measurements via the motor speed. We measure *ω* as before, using back-focal-plane interferometry [Svoboda et al., 1993] and a polystyrene bead attached to a short filament stub (see *Materials and Methods*) [Bai et al., 2010, Krasnopeeva et al., 2018, Rosko et al., 2017, Ryu et al., 2000].

Fig. 5A shows simultaneous measurements of ThT intensity and normalized motor speed during dye equilibration in MM9 + glucose. The motor speed decreases during ThT equilibration and BFM stops at the point of *I*_*in*_ decrease. Furthermore, BFM does not resume spinning even as *I*_*in*_ further increases, suggesting that the second ThT intensity increase that culminates in a plateau, is not driven by *V*_*m*_. To confirm the result, during ThT equilibration, we supplemented the medium with propidium iodide (PI). PI permeates bacterial membrane that lost its integrity and significantly enhances its quantum yield upon binding to DNA, which is commonly interpreted as an indication of cell death [Krämer et al., 2016, Lopez-Amoros et al., 1995]. We found that the cells stained with PI although ThT intracellular concentration remained high, Fig. 5B. In addition, at the time point of *I*_*in*_ decrease cellular volume suddenly increases, and cytoplasmically expressed fluorescent protein mCherry-mKate2 hybrid (referred to as mKate2 for brevity) [Lord, 2014] starts leaking out of the cell, Fig. 5C, SI Video 2.

**Figure 5.**
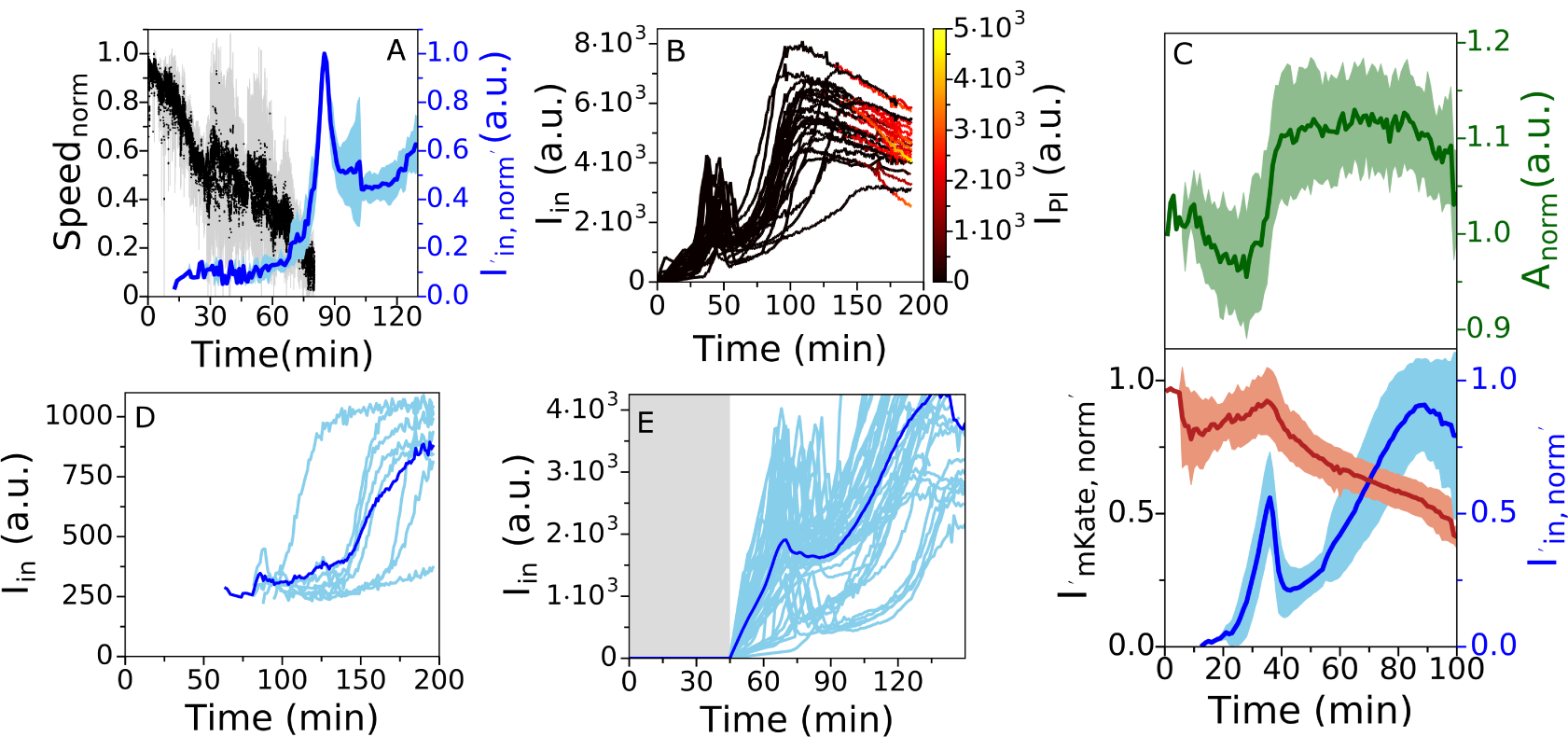
(A) Average traces of ThT fluorescence (in blue) and motor speed (in black) simultaneously measured in 5 individual cells (individual cell traces are given in Fig. SI6). The shaded areas show the standard deviation and the motor speed has been normalized to the initial value as described in *Materials and Methods*. (B) ThT (y axes) and PI (colourmap) equilibration profile in LB. 25 individual traces are given. (C) Average of ThT (in blue), mKate2 (in red) fluorescence and cell area (in green) simultaneously measured in 12 individual cells from 3 independent experiments. The shaded areas show the standard deviation. (D) Equilibration profile of 1 *µ*M ThT in LB. 8 single cell traces and average trace are given in cyan and blue, respectively. (E) Equilibration profile of 10 *µ*M ThT in LB in the absence (gray shaded area) and presence of epifluorescent illumination (light area). The dye was flown in the flow cell for the whole length of the experiment, imaging conditions in the light area were the same as in Figure 3 and 5D. 44 cells from 8 independent experiments are given.

These results are in contradiction with our estimate of dye working concentration, and we wondered, based on Fig. 2C, if the changes in *P*_*Dye*_ could be the explanation. The cell culture in Fig. 3 was briefly exposed to light at 600 nm every 7.5 min, whereas cells in our flow-cell were exposed to light of 435 nm every minute for the purpose of imaging the ThT dye. We have previously reported loss of *V*_*m*_ and PMF due to light-induced decrease of *E. coli* membrane’s resistance at effective powers higher than *∼*17 mW/cm^2^, and for a combination of 395 and 475 nm wavelengths [Krasnopeeva et al., 2018]. Light damage is wavelength dependent [Vermeulen et al., 2008], and we therefore characterized the light damage caused by our imaging conditions, i.e. 435 nm wavelength and effective power of P_eff_ *∼*11 mW/cm^2^. Fig. SI7 shows a decrease in BFM motors’ speed, and thus cell’s PMF. However, the PMF is not fully lost, indicating that the loss of PMF observed in Fig. 5A is likely caused by the combination of light induced increase in *P*_*Dye*_ and exposure to 10 *µ*M ThT.

To prove it, we exposed the bacteria to 10 *µ*M ThT in LB as before, but this time we observe the cells under bright-field illumination for 45 min, at which point we turn on the 435 nm light used for epifluorescent imaging of ThT. Fig. 5E shows that after 45 min cells not exposed to 435 nm light did not take up ThT, in contrast to Fig.3D where cells exposed to 435 nm light from the start, took up ThT in the first 30 min.

Actively changing membrane permeability has been used to facilitate loading of Nernstian sensors [Lo et al., 2007], and Fig. 5E shows that this can change the dye into an actuator because it can influence *V*_*m*_. Our mathematical model predicts that if a given concentration of the dye is lowering *V*_*m*_, an even lower concentration of the dye will result in a change of *τ*_*eq*_ (Fig. SI3). In agreement, Fig. 5D shows that for 1 *µ*M concentration of ThT *τ*_*eq*_ lasted longer then with 10 *µ*M. Thus, 10 *µ*M ThT in LB, under 435 nm light, affects *V*_*m*_. Assessing the suitability of the dye working concentration by confirming that a lower dye concentration does not alter *τ*_*eq*_ is a suitable additional control we propose, especially if *P*_*Dye*_ is being altered as part of the experiments.

We note that in our plate reader experiments (Fig. 2) we observed the effect of the dye (above 10*µ*M) on cell growth, while in our microscopy experiments, in the absence of light damage, ThT does not permeate WT cells. We thus wanted to confirm that at higher concentrations ThT permeates the cells and for the purpose we imaged the cells from the wells at representative ThT concentrations in MM9+glucose (10, 50 and 100 *µ*M) and MM9+glycerol (10 *µ*M). We found that in MM9 glucose cell brightness increases with the extracellular dye concentration and that in MM9+glycerol, at 10*µ*M, ThT signal from the cells is overall greater that in glucose (Fig. SI8).

Having identified the mechanisms behind the shape of the ThT loading curve we observed in Fig. 3 we should now be able to reproduce it with our mathematical model. Based on Fig 5A we assume that *V*_*m*_ decays exponentially immediately after addition of the dye [Krasnopeeva et al., 2018]: 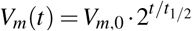 where *t*_1*/*2_ is the time at which voltage is half that of *V*_*m*,0_. The dynamics of dye entry are then modelled by Eyring’s rate law Eq. (9) taking into account *V*_*m*,0_, *t*_1*/*2_ and *P*_*Dye*_ (see *Supplementary Information* for further details on the model). Fig. SI9 shows that the model reproduces the peak in [*Dye*]_*in*_ observed in Fig. 3. Immediately upon addition, the positively charged dye moves inwards because its extracellular concentration is higher than the intracellular and the cell is negatively polarized. Thus, [*Dye*]_*in*_ increases and becomes greater than [*Dye*]_*out*_ until the electrochemical potential reaches Δ*G*_*Dye*_ = 0 (at the peak). As the *V*_*m*_ decays and since [*Dye*]_*in*_ *>* [*Dye*]_*out*_, the dye now starts moving outwards and its intracellular concentration decreases. The time at which the peak occurs as well as its intensity depend on *P*_*Dye*_ as follows: *(i)* the time of the peak decreases with increasing *P*_*Dye*_ and increases with increasing *t*_1*/*2_, and *(ii)* the intensity of the peak increases with increasing *P*_*Dye*_ and *t*_1*/*2_, Fig SI9 C and D.

The dye still equilibrates according to equation (2), but this is achieved transiently at the time of the peak, which is the time point at which *V*_*m*_ can be calculated from equation (2). Since the *V*_*m*_ varies during the course of the experiment, the *V*_*m*_ measured at the peak is not equal to the *V*_*m*,0_. Nonetheless, if we measure *V*_*m*_(*t*)*/V*_*m*,0_ as well as calculate the *V*_*m*_ at the time of the peak using Eq. (2), in principle we can estimate *V*_*m*,0_ as well. Thus, charged dyes can be used to estimate initial *V*_*m*_ even in conditions where they act as actuators and collapse *V*_*m*_, if the dynamical shape of the *V*_*m*_ loss is known.

## DISCUSSION

Nernstian probes are a popular choice for estimating bacterial *V*_*m*_, because their concentration directly depends on *V*_*m*_ according to the Nernst equation. Despite the wide usage, the probes are often not sufficiently calibrated before use in different conditions. Here we present a mathematical model that shows trade-offs between requirements imposed on the dye: sufficient signal-to-noise ratio, sufficiently short dye equilibration time and minimum effect on the cells’ physiology. Based on the model results we propose a general work-flow for the characterization of Nernstian dye candidates (Fig. 6), and demonstrate it on a newly suggested, and previously uncharacterized in *E. coli*, fluorescent dye Thioflavin T. We find that the suitability of a candidate Nernstian dye is context dependent, and reveal that the candidate dyes can be substrates for bacterial drug-export systems. We believe our work-flow is sufficiently simple and general to provide a common standard for benchmarking the cationic dye behavior and thus improve the robustness of *V*_*m*_ measurements.

**Figure 6.**
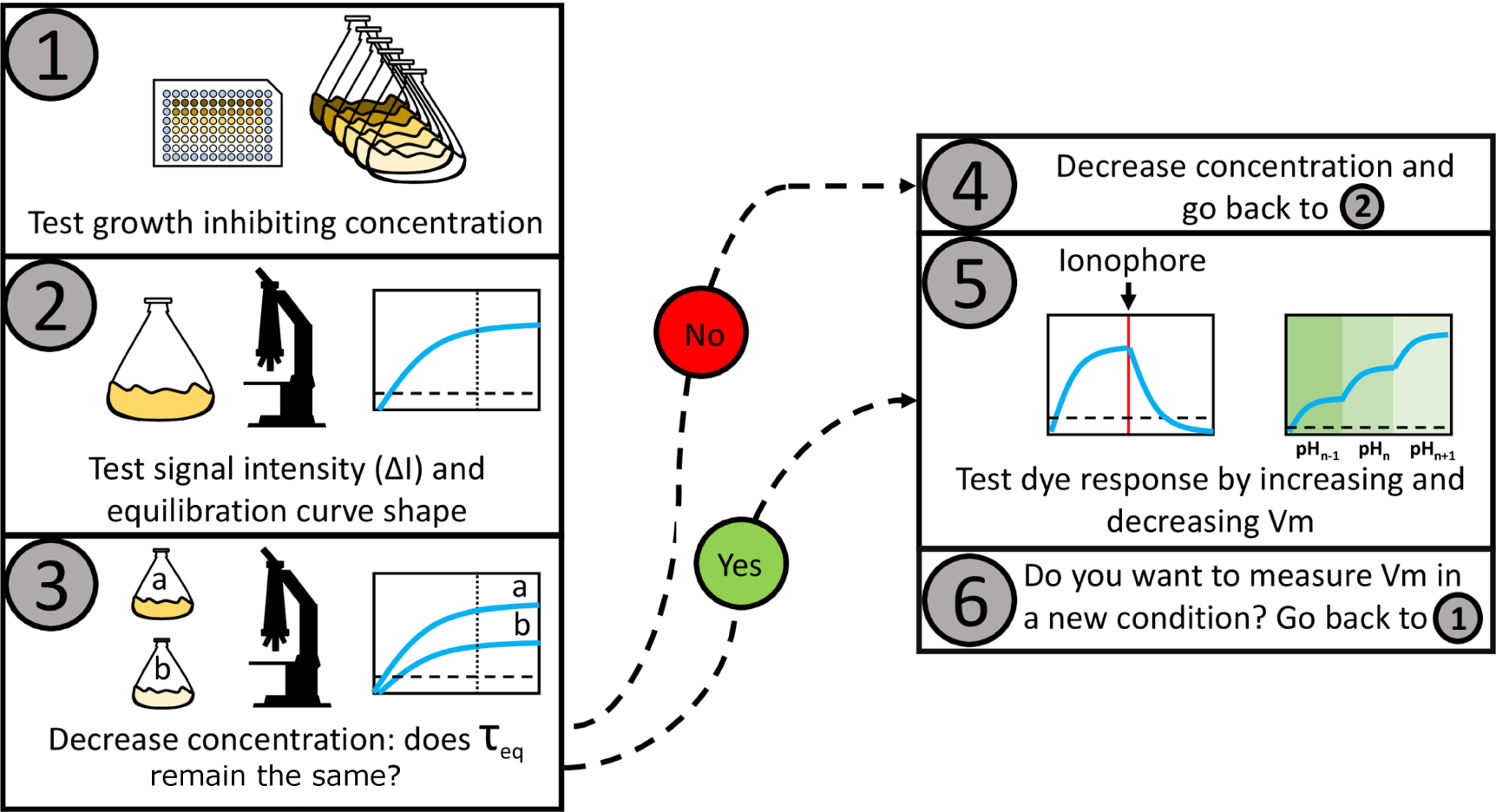
Proposed work-flow for characterizing the Nernstian behavior of a candidate cationic membrane voltage dye. The working concentration is estimated in steps 1 to 4, we define it as the maximum dye concentration that does not affect membrane voltage and that yields sufficient amount of signal. The MNC is estimated. (2) The MNC is tested for sufficient signal intensity. (3) The effect of the dye on *V*_*m*_ is determined by measuring *τ*_*eq*_ at different below-MNC concentrations. (4) Different *τ*_*eq*_ for different dye concentrations indicate that the probe is altering *V*_*m*_ and the working concentration should be reduced and the protocol resumed from (2). Equal *τ*_*eq*_ indicates that the probe is not altering *V*_*m*_. (5) Common procedures to test Nernstian dye responses can then be applied, such as the introduction of a ionophore that neutralizes *V*_*m*_, or changes in external pH that induce changes in *V*_*m*_ [Lo et al., 2007]. (6) Because the effects depend on the physiological state of the cell, the procedure should be repeated for every experimental condition.

We found as well that imaging conditions can affect how well the dye crosses bacterial membrane, which suggest that on its own *E. coli* membrane exhibits low permeability to this Nernstian sensor, or that its export by membrane efflux pumps is substantial. The observation is consistent with previous results that needed to permeabilize the membrane by EDTA to achieve experimentally reasonable loading times [Lo et al., 2007]. However, changing the membrane permeability is often not dye specific, and can lead to changes in the overall *V*_*m*_ of the cell. Our model suggests that if the chosen dye working concentration is not affecting *V*_*m*_, lower dye concentration should leave *τ*_*eq*_ unchanged. The finding offers a simple test to confirm the suitability of the chosen dye working concentration, which should be performed in each environmental and physiological condition.

As a final remark, the fact that in the absence of illumination of shorter wavelengths, i.e. in our plate reader experiments, ΔTolC mutant is affected significantly more by the dye than the wild type suggest that, in the rich defined medium such as MM9 + glucose and amino acids, low cytoplasmic dye concentration is not just a consequence of low membrane permeability to the dye, but also of the active efflux. Then, in a condition where membrane permeability is affected, leading to the PMF loss, which is in turn the energy source for efflux pumps, the cytoplasmic concentration of the dye is no longer in accordance with equation (2) and could result in interesting dynamics such as in Fig. 5A, where we observe significant accumulation of the dye only after the PMF drops to a certain threshold value.

## AUTHOR CONTRIBUTIONS

LM, GT, CJL, BF and TP conceived the experiments and the computational work. LM, TT, YP and YL performed experiments. GT performed computational work. LM analysed experimental data, and LM, GT, CJL, BF and TP interpreted the results and wrote the manuscript.

### ACKNOWLEDGEMENTS

We thank Dario Miroli for help with image analysis, Nathan Lord and Sebastian Jaramillo-Riveri for donating us the construct containing the hybrid mCherry-mKate2 sequence and Angela Dawson for retrieving the ΔTolC mutant from the Keio collection. This work was financially supported by the Cunningham Trust scholarship ACC/KWF/PhD1 to TP and LM, the National Natural Science Foundation of China under grants No. 31722003 and No. 31770925 to FB, the Ministry of Science and Technology, Republic of China, under contract No. MOST-107-2112-M-008-025-MY3 to CJL and Human Frontiers Program grant RGP0041/2015 to TP, FB and CJL. TP acknowledges the support of UK Research Councils Synthetic Biology for Growth programme and is a member of the BBSRC/EPSRC/MRC funded Synthetic Biology Research Centre (BB/M018040/1).

